# Uniform distribution of photosystems in dark-adapted *Synechocystis* cells

**DOI:** 10.1101/2025.10.29.685299

**Authors:** Peter R. Bos, Elio Langlois – Legrand, Emilie Wientjes

## Abstract

Photosynthesis in cyanobacteria relies on light capture by photosystem I (PSI), photosystem II (PSII) and the phycobilisome (PBS). While these complexes are thought to be intermixed within the thylakoid membrane, there is also evidence for PSI-enriched and PSII-PBS-enriched microdomains, and their spatial organization is still debated. This organization may further depend on environmental conditions. To study it, a range of methods are available. Cryo-electron tomography offers the highest resolution but is limited in throughput, while super-resolution fluorescence techniques such as Airyscan and structured illumination microscopy provide improved resolution but cannot resolve individual thylakoid membranes. To expand this toolbox, we applied cryo-Expansion Microscopy (cryo-ExM) to dark-adapted *Synechocystis* sp. PCC 6803 cells. Cells were cryofixed, rehydrated at room temperature and physically expanded in a swellable hydrogel. By expanding cells 5.5-fold, we resolved individual thylakoid membranes in intact cells using confocal microscopy. Immunostaining further allowed simultaneous localization of PSI, PSII and PBS within the expanded thylakoid network. Quantitative analysis of fluorescence covariance revealed a high degree of colocalization among PSI, PSII and PBS, providing no evidence for microdomains under dark-adapted conditions. PBS was excluded only from the neck region between dividing cells, while PSI, PSII and PBS were otherwise distributed throughout the thylakoid membrane. Together, these results establish cryo-ExM as a powerful method for visualizing individual cyanobacterial thylakoid membranes and mapping the distribution of key photosynthetic complexes, thereby complementing existing approaches for dissecting the spatial organization of photosynthesis.

## Introduction

The photosynthetic membrane of cyanobacteria contains Photosystem I (PSI), Photosystem II (PSII), cytochrome *b6f*, and ATP synthase as the major membrane-embedded protein complexes that drive photosynthesis (Rexroth et al., 2017). Together, these complexes perform linear and cyclic electron transport to generate NADPH and ATP, which are subsequently used in CO_2_ fixation via the Calvin–Benson cycle (Lea-Smith et al., 2016). In contrast to higher plants, where the PSI/PSII ratio is typically below one, cyanobacteria like *Synechocystis* sp. PCC 6803 (from now in *Synechocystis*) and S*ynechococcus elongtus*, exhibit a higher PSI/PSII ratio ranging from two to five, reflecting a different balance in light harvesting and energy distribution (Casella et al., 2017; MacGregor-Chatwin et al., 2017; Moore & Vermaas, 2024). Another fundamental difference is that, in cyanobacteria, photosynthesis and respiration both occur in the thylakoid membrane system, whereas in plants these processes are spatially separated (Lea-Smith et al., 2016; Mullineaux, 2005). This shared localization requires a different organizational logic of the membrane and energy-transfer network.

Instead of the membrane-embedded light-harvesting complexes of plants, cyanobacteria generally employ phycobilisomes (PBSs) as their main antenna system (Adir et al., 2020). PBSs are large supramolecular complexes, up to ∼60 nm in size, attached to the photosystems or to the thylakoid membrane. PBSs typically consist of a core of allophycocyanin, with rod-like extensions primarily containing phycocyanin and, in some species, additional pigments such as phycoerythrin or phycoerythrocyanin. PBS size and composition are dynamically regulated by light conditions, allowing flexible adjustment of energy transfer (Adir et al., 2020; Kehoe & Gutu, 2006). Importantly, excitation energy harvested by PBSs can be transferred to both PSI and PSII, depending on the physiological state of the cell, although the extend of PBS-PSI transfer remains under debate (Akhtar et al., 2022; Choubeh et al., 2018; Chukhutsina et al., 2015; van Stokkum et al., 2025). In addition, spillover of excitation energy between PSII and PSI has been suggested to occur in cyanobacteria (Ma et al., 2008; Ueno et al., 2017).

In higher plants, spillover between PSII and PSI is minimized by the strict spatial separation of photosystems into distinct thylakoid regions: PSII is enriched in grana stacks, while PSI is located primarily in stroma lamellae (Andersson & Anderson, 1980; Dekker & Boekema, 2005). This heterogeneity is critical for efficient regulation, as disruption of grana stacking leads to growth defects due to impaired light-energy tuning (Armbruster et al., 2013; Pribil et al., 2018; Trissl & Wilhelm, 1993). Cyanobacteria, by contrast, lack grana and do not display such strict partitioning of PSI and PSII, raising the question of how they maintain photosynthetic efficiency without this structural separation (Mullineaux, 1992, 2005; Mullineaux & Holzwarth, 1993).

An increasing number of studies suggest that cyanobacteria may nevertheless exhibit a form of heterogeneity through the formation of ‘microdomains’. These microdomains are regions enriched in either PSI or PSII of 0.5-1.5 µm in size, but without the strict segregation observed in plants (Strašková et al., 2019). Evidence for such domains has been reported based on *in vivo* and *in vitro* studies, using biochemical fractionation, atomic force microscopy (AFM), confocal microscopy, or super-resolution imaging (Agarwal et al., 2010; Canonico et al., 2020; Casella et al., 2017; Collins et al., 2012; Huokko et al., 2021; Kaňa et al., 2023, 2024; Sherman et al., 1994; Strašková et al., 2019; Vermaas et al., 2008). However, several questions remain regarding the precise nature of these domains and the mechanisms underlying their formation. Firstly, it is still unclear whether microdomains form passively due to membrane dynamics or actively through regulatory processes. Secondly, the effect of isolation is unclear for the *in vitro* studies or techniques that require isolated membranes. Thirdly, PSI-enriched regions in *S. elongatus* have been observed in either central thylakoids or outer thylakoids, depending on the study (Sherman et al., 1994; Vermaas et al., 2008). Lastly, earlier studies with EM and confocal microscopy in *Synechocystis* did not find evidence for a heterogeneous distribution of PSI and PSII, thereby raising the question how lower-resolution imaging techniques were able to distinguish those domains in other studies (Collins et al., 2012; Liberton et al., 2013; Van De Meene et al., 2006).

Here, we applied cryo-Expansion Microscopy (cryo-ExM) to dark-adapted *Synechocystis* sp. PCC 6803 to directly visualize its cellular structure and, in particular, the organization of the photosynthetic membrane. Cryo-ExM combines rapid fixation in liquid ethane at 77K with ExM, in which a sample is physically expanded using a swellable hydrogel. After freezing and fixation, the sample is brought up to room temperature, anchored and embedded in a swellable hydrogel. The technique has been shown to better preserve cytoskeletal elements than conventional chemical fixation (Laporte et al., 2022). Moreover, Cryo-ExM has been successfully applied to whole *Chlamydomonas reinhardtii* cells, including immunolabeling of PSII subunits (Klena et al., 2023; Laporte et al., 2022). In our study we used antibodies against PsbA (PSII), PsaC (PSI), and C-phycocyanin (PBS) to localize the three key components of the light-harvesting machinery. With the ∼5.5-fold resolution improvement compared to confocal microscopy, we did not detect evidence for microdomains. Instead, we observed almost complete covariance between PSI, PSII, and PBS signals, and their peak intensities overlapped, suggesting a largely homogeneous distribution across the thylakoid layers of dark-adapted *Synechocystis*.

## Material and Methods

### Cyanobacterial growth

#### Synechocystis sp

*PCC 6803* and *Synechococcus elongatus PCC 7942* were grown in 100 ml of BG-11 medium at 30°C under white light illumination at 30 μmol photons m^-2^s^-1^ in 250 ml flasks shaken at 160 rpm, containing a stirrer. The original cultures were grown on BG-11 with 1.5% agar plates under the same conditions.

### Fixation

#### Cryofixation

Cyanobacterial cultures were grown for two weeks after inoculation from plate. Cells were washed twice by centrifuging at 800 g for 5 min and resuspension in PBS. Then, a drop of the cell culture was placed on a 12 mm diameter cover slip, that was coated with poly L-lysine and allowed to sediment for at least 10 min in the dark. Next, we used the Vitrobot Mark IV System by Thermofisher to plunge freeze the slides in liquid ethane according to the cryo-ExM protocol (Klena et al., 2023; Laporte et al., 2022). After plunging, the slides were transferred to liquid nitrogen frozen acetone containing 0.1% PFA and 0.02% GA and left overnight on dry ice, where the acetone melts again. The next morning, the tubes were allowed to slowly come to room temperature by removing most of the dry ice and allowing the rest to evaporate. Then samples were rehydrated by sequential incubation for 5 min in solutions of 100% ethanol, 95% ethanol (2x), 70% ethanol, 50% ethanol, Milli-Q water, and PBS. The samples were then anchored using a 1.4% paraformaldehyde, 2.0% acrylamide solution in PBS and incubated for 3h at 37°C in a humid environment.

### Expansion microscopy

Next, we applied ExM as before (Berentsen et al., 2025; Bos et al., 2024), with adaptations in the gelation chamber design and gel denaturation. In short, a gel composition of 23% sodium acrylate, 8.9% acrylamide, 0.09% N,N′-methylenebisacrylamide in phosphate buffered saline was made and stored at least one night at -20 ºC. To this gel we added a 10% TEMED solution and a 10 % APS solution to a final concentration of 0.1% to start polymerization. The slide with the fixed and anchored cells was carefully blotted dry without touching the cells. A 25 μL drop of gel solution was placed on a piece of parafilm on ice and the slide containing the cells was placed on the drop with the cells facing the solution. Polymerization was allowed to occur for 5 minutes on ice after which the gels were moved to a humid chamber and polymerized further at 37 ºC for 90 minutes.

The gels were removed from the slides with a scalpel and denatured at 95 ºC for 90 minutes in a denaturation buffer (200 mM SDS, 200 mM NaCl and 50 mM TRIS base, pH 9.0). The denatured gels were washed at least three times in ultrapure water and left overnight in water to reach full expansion and wash away remaining SDS. A piece of the central part of the gel was cut out and used for staining. A more detailed explanation of this method can be found in Klena’s protocol article (Klena et al., 2023).

### Antibody and all-protein staining

For the antibody staining, we used rabbit polyclonal antibodies targeted against PsbA at 1:200 concentration (Product no: AS05 084) and PsaC at 1:200 (Product no: AS10 939) both purchased from Agrisera (Agrisera AB, Sweden). PsbA was reported to have little off target binding proteins, while one extra band was reported fot anti-PsaC in *Synechococcus*, as reported on the manufacturer’s website. Rabbit anti C-Phycocyanin antibody was applied at 1:50 (AbBy Fluor® 594) and purchased from Antibody-online (No. ABIN2802262). This antibody reported little off-target effects (Geh et al., 2015), as was reported for other anti-C-phycocyanin antibodies in *Synechocystis* (CpcA / Anti-C-phycocyanin alpha subunit Antibody for phytoab, USA).

Anti-PsbA was conjugated with NHS-ATTO594 or NHS-ATTO647 (ATTO-TEC GmbH, Art. Nr.: AD 594 or 647) by adding 0.2μL of ATTO dye to 15μL of antibody. The reaction was allowed to proceed for 1h, and excess dye was removed by using a desalting column (Zeba spin 7K MWCO Column). The column was washed 4 times with PBS prior to loading the labelled protein solution. After loading, the column was centrifuged (1500 xg, 1 minute) and the eluted fraction stored in the freezer. The eluted fraction was typically larger than the initial 15 μL of protein, which was taken into account when determining the antibody dilution for staining. Each antibody staining was diluted in a solution of 0.2% bovine serum albumin in PBS. Once deposited on the gels, the gels were incubated at 37 °C for 3h and washed three times with PBS with 0.1% tween for 10 min per wash.

Multiplexing was achieved by first staining with anti-PsaC for 3h at 37°C followed by two washing steps with PBS-Tween 0.1%. Then its secondary antibody (Goat anti-Rabbit IgG (H+L) Cross-Adsorbed Secondary Antibody, Alexa Fluor™ 488 (AB_143165), ThermoFisher, or Nano-Secondary® anti-rabbit IgG, recombinant VHH, Alexa Fluor® 488 (CTK0102), ChromoTek GmbH & Proteintech Germany) was applied and left for 1.5 hr at 37°C. After that, Anti-C-phycocyanin and anti-PsbA were used simultaneously to avoid extra washing steps with the 3 hours incubation time and 37°C, followed by 2 washing steps with PBS-0.1% Tween.

#### All-protein staining

To create an all-protein staining of the sample, we applied ATTO dyes with an NHS-ester tag. The NHS-ester binds lysine and therefore creates a lysine or protein density staining of the sample (M’Saad & Bewersdorf, 2020). Depending on the other fluorophores present, we used 20 μg/mL N-hydroxysuccinimide (NHS) ester-ATTO488 (ATTO-TEC GmbH, Art. Nr.: AD 488), ester-ATTO594 (ATTO-TEC GmbH, Art. Nr.: AD 594) and ester-ATTO647 (ATTO-TEC GmbH, Art. Nr.: AD 647) in 0.1 M NaHCO3, pH 8.3 with 100μL per gel in 8 well plate and then incubated for 1.5 h (M’Saad & Bewersdorf, 2020).

### Image acquisition

Cut and stained gels were placed in an 8-wells plate and left overnight to reduce drift of the gel during imaging. At low laser power, drift was usually negligible, but when higher laser powers were required, the drift became too large to be corrected for. In those instances, gels were placed in a PLL coated 8-wells plate.

Samples were imaged using Leica Stellaris confocal microscope equipped with an HC PL APO CS2 86×/1.20 water objective. This system automatically sets the system optimized settings for all dyes and imaging modes. ATTO-488 and Alexa-488 were excited at 501 nm and emission was recorded from 506-579 nm, ATTO 594 and Alexa 594 were excited at 590 nm and emission recorded from 595-648 nm and ATTO647 was excited at 646 nm and emission recorded from 660-830 nm. Counting mode was used for all channels and 4-8 line accumulations, depending on the intensity of the dyes. If ATTO594 and ATTO647 were used simultaneously, crosstalk was prevented by line sequential scanning. ATTO 488 and ATTO 647 were detected in one sequence and ATTO594 in another. System optimized settings were used for the voxel size, which was typically around 83×83×356 nm in XYZ. 3D models were created from z-stacks in the Leica LAS X software.

### Image analysis

Image analysis was performed in FIJI (Schindelin et al., 2012) and Python3.9.

#### Pearsons’s coefficient

Images of expanded *Synechocystis* cells, stained with anti-PSI, PSII and Phycocyanin antibodies were selected for colocalization analysis. Slides with the highest intensity in z-stacks were selected by hand. They were analyzed in Python3.9 with the package Scikit-Image (Van der Walt et al., 2014) in a custom written script. We blocked out all pixels that were not part of the cyanobacterium by applying a gaussian blur on the image of 7 pixels, followed by an Otsu filter (SI figure 3B). This mask was multiplied with the original image (SI figure 3A and C), and an Otsu filter was applied again on the resulting image (SI figure 3D). The Pearsons coefficient is a measure for the linearity between the pixel intensities of two images. This value was determined on two different channels, treated as SI figure 3C (Schober et al., 2018; Sedgwick, 2012).

#### Peak position

A custom written script was used to determine the peak position of the three channels (SI figure 4). A gaussian blur and Otsu filter were applied and the X and Y position with the highest number of pixels above the threshold were selected. The intensity profiles of all channels were taken and smoothed with a Satvitsky Golay filter with a window size of 25 and a 4^th^ order polynomial. Peaks that were at least 0.3*max intensity, 20 pixels apart and a peak prominence larger than 1.0, were selected. If those peaks in all channels were less than 15 pixels apart, the difference was determined between peak position and the mean peak position. The difference in pixel position was calculated to pre-expanded dimensions by multiplying by the pixel size (83 nm) and divided with the expansion factor (5.5).

## Results

To identify the localization of the key protein complexes in photosynthesis in *Synechocystis*, we applied cryo Expansion Microscopy (cryo-ExM). We first cryo-fixed the cells (Klena et al., 2023; Laporte et al., 2022) in liquid ethane and after rehydration of the sample, embedded it in a swellable hydrogel. The samples were denaturated, expanded and stained with an all-protein NHS ester dye. An expansion factor of 5.5 was sufficient to resolve the different thylakoid layers (figure 1). Next to the thylakoid membranes, the carboxysomes were clearly visible as bright spheres in the center of the cell. An intensity plot showed the thylakoid membranes to be roughly 91 ± 20 nm apart, but on occasions we could distinguish layers that were less than 60 nm apart in pre-expanded dimensions. This separation distance is somewhat larger than the 78 nm reported before in *Synechococcus elongatus* (Huokko et al., 2021) and considerably larger than the 62 nm reported for *Synechocystis* with neutron scattering (Liberton et al., 2013). However, with our technique we are unable to distinguish thylakoid layers that are closer than 60 nm, and might therefore overestimate the average thylakoid distance.

**Figure 1.**
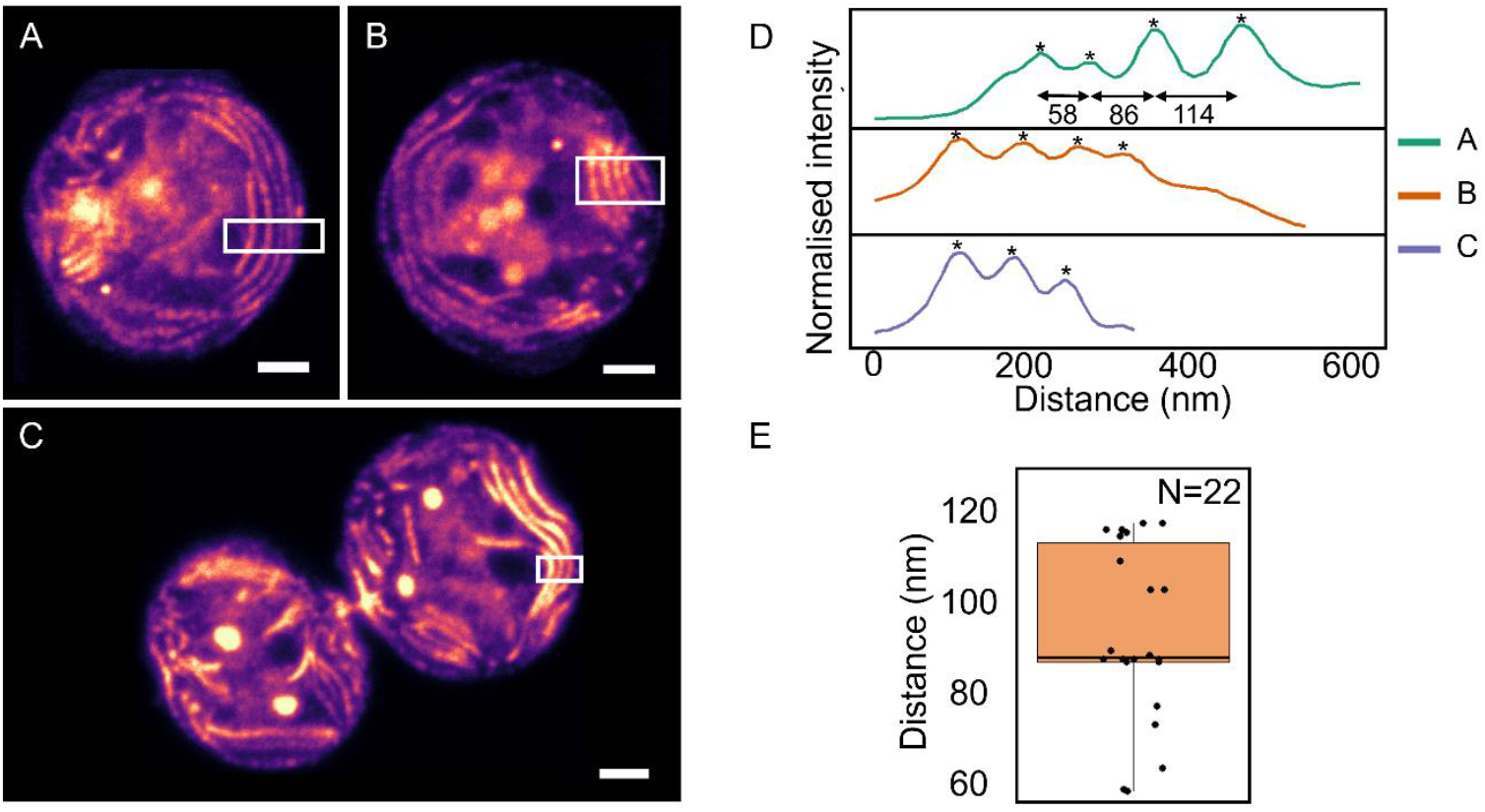
A, B, C) Micrographs of expanded *Synechocystis* cell, stained with an all-protein staining. Scale bars represent 1 µm in pre-expanded dimension. In white boxes, regions from which the profiles are plotted in D. Peak positions of the thylakoid membrane are indicated by asterisks and for A the differences between the peaks are indicated in nm, pre-expanded dimensions. E Peak positions from D and other images, plotted in a boxplot. 95^th^ percentile and 5^th^ percentile are indicated by whiskers, the box represents 25^th^ to 75^th^ percentile and the line in the box the median.

We localized PSI, PSII and the PBS using antibodies against PsaC, PsbA and phycocyanin, respectively. Overlay of the all-protein NHS-ester staining with the antibody labeling demonstrated that all three antibodies specifically localized to the thylakoid membrane (figure 2A, B and C). For example, the carboxysomes, which are protein dense regions (MacCready & Vecchiarelli, 2021), were not visible in the antibody channels. Negative control with only secondary antibody did not show intensity (SI Figure 2). Next, we localized the three protein complexes together in the same sample. Merging of the micrographs indicated almost complete overlap of the antibody signals (figure 2D, E and F).

**Figure 2.**
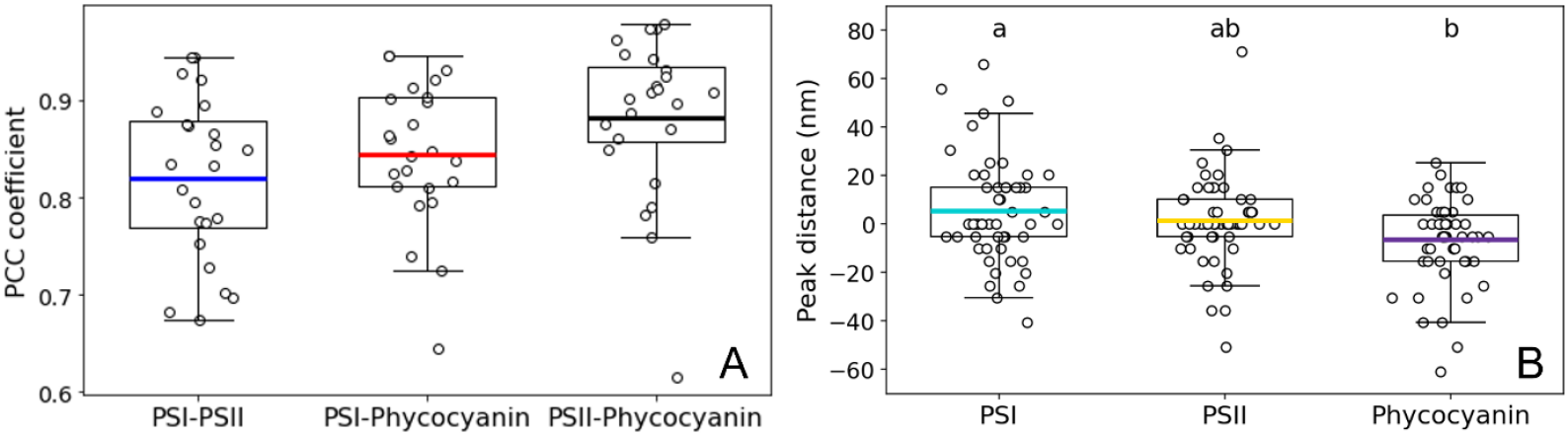
Measures for colocalization of the three antibodies. A) PCC for all three pairs of antibodies. B) Difference in peak position of the individual antibodies from the mean peak position. A negative position means localization away from the center than the mean, ie outside the cell and a positive position represents localization inside the cell. Each dot represents an individual measurement and the line represents the mean. The box shows 25^th^ and 75^th^ percentile and the whiskers 5^th^ and 95^th^ percentile. Examples of the image analysis are presented in SI figure 3 and 4.

The Pearsons Colocalization Coefficient (PCC) was determined between the pairs of antibodies. The PCC determined covariance between channels, with -1 indicating perfect anticorrelation and 1 indicating perfect correlation between pixel intensity. For all three pairs (PSI-PSII, PSI-phycocyanin and PSII-phycocyanin) we find a PCC higher than 0.8, indicating a high degree of correlation in their pixel intensities (figure 3A). No significant differences were found between the pairs.

**Figure 3.**
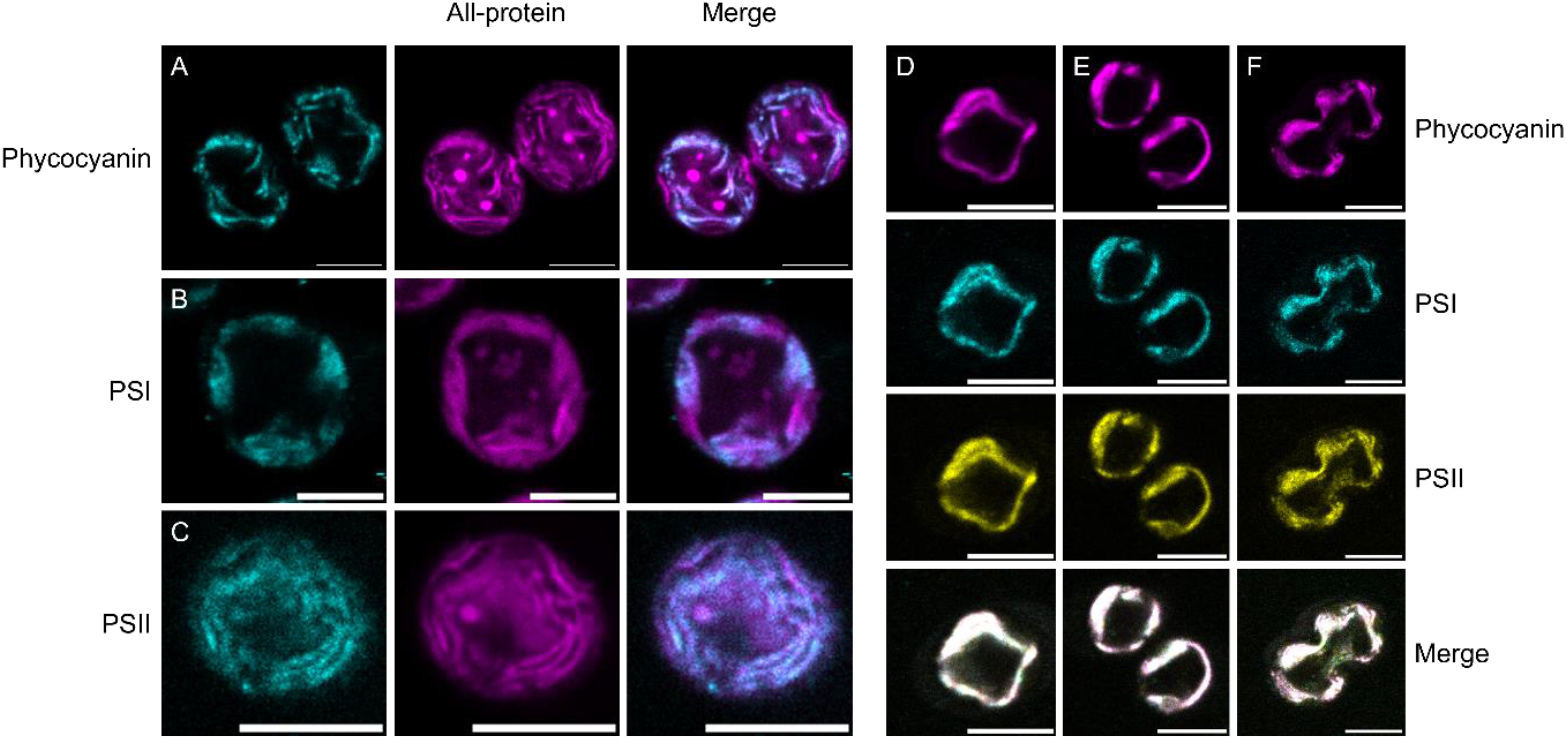
A, B and C) Example micrographs of expanded *Synechocystis* cells, immunolabelled (cyan) and pan-protein stained (magenta). D, E and F) Immunolabelled expanded *Synechocystis* cells. The cells are simultaneously stained with anti-PsaA, anti-PsbA and anti-phycocyanin. Scale bars represent 1 µm in pre-expanded dimensions.

Next, we addressed the question of preferential PSI localization either towards the center or the periphery cyanobacterial cells (Collins et al., 2012; Sherman et al., 1994; Strašková et al., 2019; Vermaas et al., 2008). Specifically, we determined the peak intensity of PSI, PSII, and PBS within individual cells (SI Figure 4) and calculated the deviation of each complex from the mean peak position (Figure 3B). This analysis revealed a small (20 nm on average), but significant difference between the peak positions of PSI and phycocyanin, with PSI concentrated slightly more toward the cell center and phycocyanin somewhat more to the periphery.

In most images, we detected almost complete colocalization of PSI, PSII and PBS, however, some exceptions were found, especially in stages of division (figure 4). Several cells had nearly completed binary fission but remained connected by a narrow tube. The connecting region contained thylakoids with PSI and PSII (figure 4A), yet it showed no detectable binding of the phycocyanin antibody. In addition, we observed one cell with an unusual morphology in which one part of the cell bulged outward, possibly representing an atypical division event. The bulging region lacked anti-phycocyanin signal but still contained PSI and PSII. In the ‘neck’ between dividing cells, the thylakoid layers were closer together than elsewhere in the cell in the all-protein stained cells (figure 1C and 2A), potentially excluding the large PBS from these parts. However, the resolution of imaging in the bulging regions was insufficient to resolve the thylakoid distance. Therefore, the reason for phycocyanin exclusion remains unknown.

**Figure 4.**
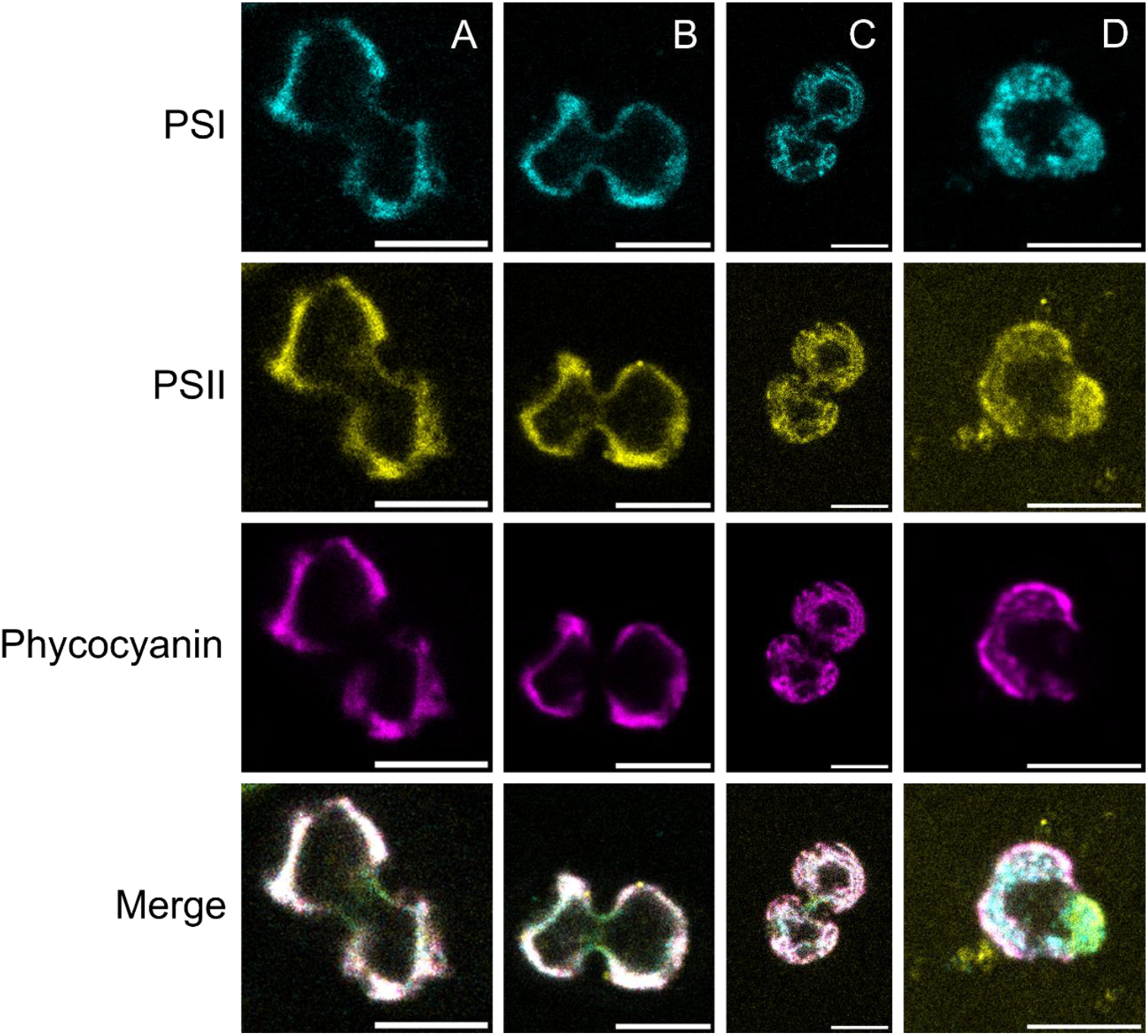
Example micrographs showing regions in which PSI and PSII colocalized but phycocyanin was not detected. For A, B and C, phycocyanin is not found in the parts linking the two cells. In D, the bulging part of the cell is devoid of anti-phycocyanin staining. PSI is shown in cyan, PSII in yellow and phycocyanin in magenta. In case of overlap, the regions turn white.

In conclusion, we detected nearly complete colocalization between PSI, PSII or PBS in the thylakoid membrane of *Synechocystis* with cryo-ExM, except for some confined regions of cells in a stage of division.

## Discussion

### Exploration of Cryo-ExM methodology

Cryo-ExM proved highly effective for *Synechocystis* cells without requiring cell wall-degrading enzymes. With an expansion factor of 5.5, thylakoid layers became clearly resolved and this factor was therefore adopted throughout our study. Attempts with chemical fixation as performed in earlier ExM studies on photosynthetic membranes (Berentsen et al., 2025; Bos et al., 2024) occasionally yielded expanded cells, but cells sometimes lost their round shape, suggesting anisotropic expansion. Moreover, immunolabeling resulted in less intense fluorescence. Results with *Synechococcus elongatus* PCC 7942yielded in some well expanded cells, but all gels also contained some unexpanded cells or cells that lost their rigid rod-like shape (SI figure 5). Moreover, PSI and PSII antibody staining were generally faint. Potentially, expansion was hindered by its more complex cell wall than *Synechocystis*, but cellulose digestion did not resolve this issue (data not shown). The approach of cryo-ExM seems promising for more cyanobacterial species, but could require optimalisation of the fixation, expansion and staining protocols.

The antibodies used, anti-PsaC, anti-PsbA, and anti-phycocyanin, all successfully localized to thylakoid membranes in *Synechocystis*, and not to other protein-dense regions such as carboxysomes. Among them, anti-phycocyanin gave the strongest signal. By contrast, anti-AtpB (ATP synthase β subunit) labeled patchily and partly bound to the cell wall (data not shown). This pattern might reflect microdomains of ATP synthase, as previously observed with eGFP labeling (Casella et al., 2017), although it could also result from imperfect recognition in our staining.

Together, these findings highlight Cryo-ExM as a promising tool for visualizing cyanobacterial cells and protein localization by immunolabeling. The method may be applicable to other cyanobacteria, with or without protocol modification. For species with a more complex cell wall, such as *Synechococcus elongatus*, incorporation of cell-wall– digesting enzymes may be necessary. Beyond antibody labeling, Cryo-ExM could be combined with genetically introduced epitopes (e.g. GFP tags recognized by anti-GFP antibodies). Such approaches open possibilities for studying protein targeting pathways, or the balance between photosynthesis and respiration, in cyanobacteria.

### Heterogeneous organization of PSI and PSII

Several studies using either fractionation, AFM, and microscopy have reported a heterogeneous separation of PSI and PSII into microdomains (Agarwal et al., 2010; Canonico et al., 2020; Casella et al., 2017; Collins et al., 2012; Huokko et al., 2021; Kaňa et al., 2023, 2024; Sherman et al., 1994; Strašková et al., 2019; Vermaas et al., 2008). In contrast to plants, this separation in cyanobacteria is not strict, but manifests itself as stable variations in ratios of photosystems. These domains have been reported to be 0.5-1.5 µm in size. By using specific light conditions, the separation and ratio differences can become more apparent (Canonico et al., 2020; Huokko et al., 2021). In the present study, however, we did not detect microdomains of PSI, PSII, or PBS in dark-adapted *Synechocystis* using immunolabeling in combination with cryo-ExM, an observation that is in line with some earlier EM and confocal microscopy studies (Collins et al., 2012; Liberton et al., 2013; Van De Meene et al., 2006). Except for a small difference in intensity peak position between PSI and phycocyanin, our analyses indicate a largely homogeneous distribution of photosystems within the thylakoid membranes of dark-adapted cells.

### Image analysis considerations

Most microscopy studies define and detect microdomains by comparing fluorescence intensity ratios across channels (Canonico et al., 2020; Kaňa et al., 2023, 2024; Strašková et al., 2019). While such an approach can be effective, e.g. for GFP expression vectors, it was not suitable for our workflow. Our extended sample preparation consisting of fixation, denaturation, gelation, and washing, can lead to loss of epitopes, and antibody labeling itself is variable. As a result, applying a uniform threshold across images was not feasible, and per-image thresholds would have introduced bias. For this reason, the Manders coefficient was not useful in our context. The Pearson correlation coefficient (PCC), by contrast, remains valid as long as intensity and signal-to-noise are sufficient. Here, we determined the PCC values only on pixels inside the object (the bacterial cell) of which at least one channel had an intensity higher than 0. The object PCC analysis in our images revealed a high degree of correlation in pixel intensity between all three antibodies, without significant differences between antibody pairs. We therefore conclude that there is no heterogeneous localization of photosynthetic protein complexes in the thylakoid membranes of dark-adapted *Synechocystis*.

Beyond covariance of pixel intensities, we also examined the peak intensity positions of antibody signals. We analyzed smoothed intensity profiles across the middle of *Synechocystis* cells (figure 3 and SI figure 4) and found that PSI localized slightly more in the middle of the cell than phycocyanin. However, this difference was not consistent in every cell, but was rather found in some cells. The average distance in peak intensity between PSI and phycocyanin was smaller than our resolution of imaging (an average difference of 20 nm vs a 60 nm resolution), and became significant due to the sample size (N=54). Moreover, the difference in intensity can be explained by the absolute intensity of the channels, where PSI antibodies usually gave the lowest fluorescence signal and phycocyanin the highest. Since signal intensity usually continued longer inside the cell than outside, and the signal to noise ratio in PSI was low, it could cause the peak of PSI to be located more to the inner part of the cell. Contrarily for phycocyanin, with a higher signal to noise ratio, the noise signal is better suppressed by the filters. Future work should test whether such profiles shift under light conditions that are known to alter PSI–PSII distribution (Canonico et al., 2020; Huokko et al., 2021).

#### Exceptions at dividing cell regions

The only regions with uneven distributions were confined areas of the thylakoid membrane. At connecting regions of dividing cells, PSI and PSII were present, but phycocyanin was absent. Similarly, in presumptive bulging regions preceding division, phycocyanin was not detected. Given the large size of PBS (∼40 nm), steric hindrance in confined thylakoid areas may exclude intact PBS complexes (Adir et al., 2020). It is possible that PBSs are remodeled into smaller subunits that remain associated with PSI or PSII, which could be tested using antibodies against PBS core or allophycocyanin.

## Acknowledgements

This work was supported by the Dutch Organization for Scientific Research (NWO) via a Vidi grant (VI. Vidi 192.042 to E.W.).

## Notes

### Competing Interest Statement

This research was supported by a grant from the Dutch Research Council.

