## Supplementary Information for "Uniform distribution of photosystems in dark-adapted *Synechocystis* cells"

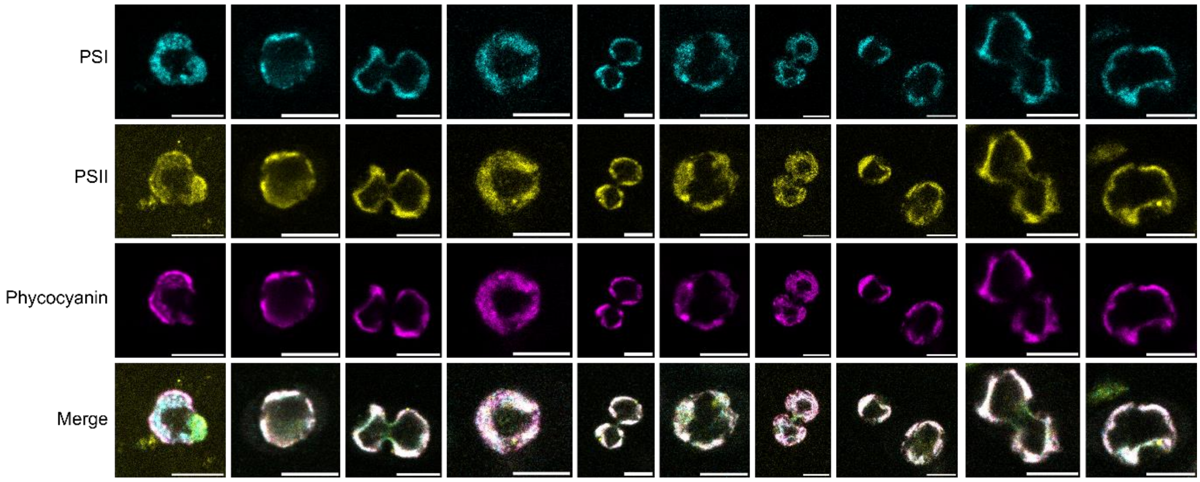


Supplemental Figure 1 Additional examples of micrographs of immunostained *Synechocystis*. Scale bars represent 1 µm in pre-expanded dimensions.

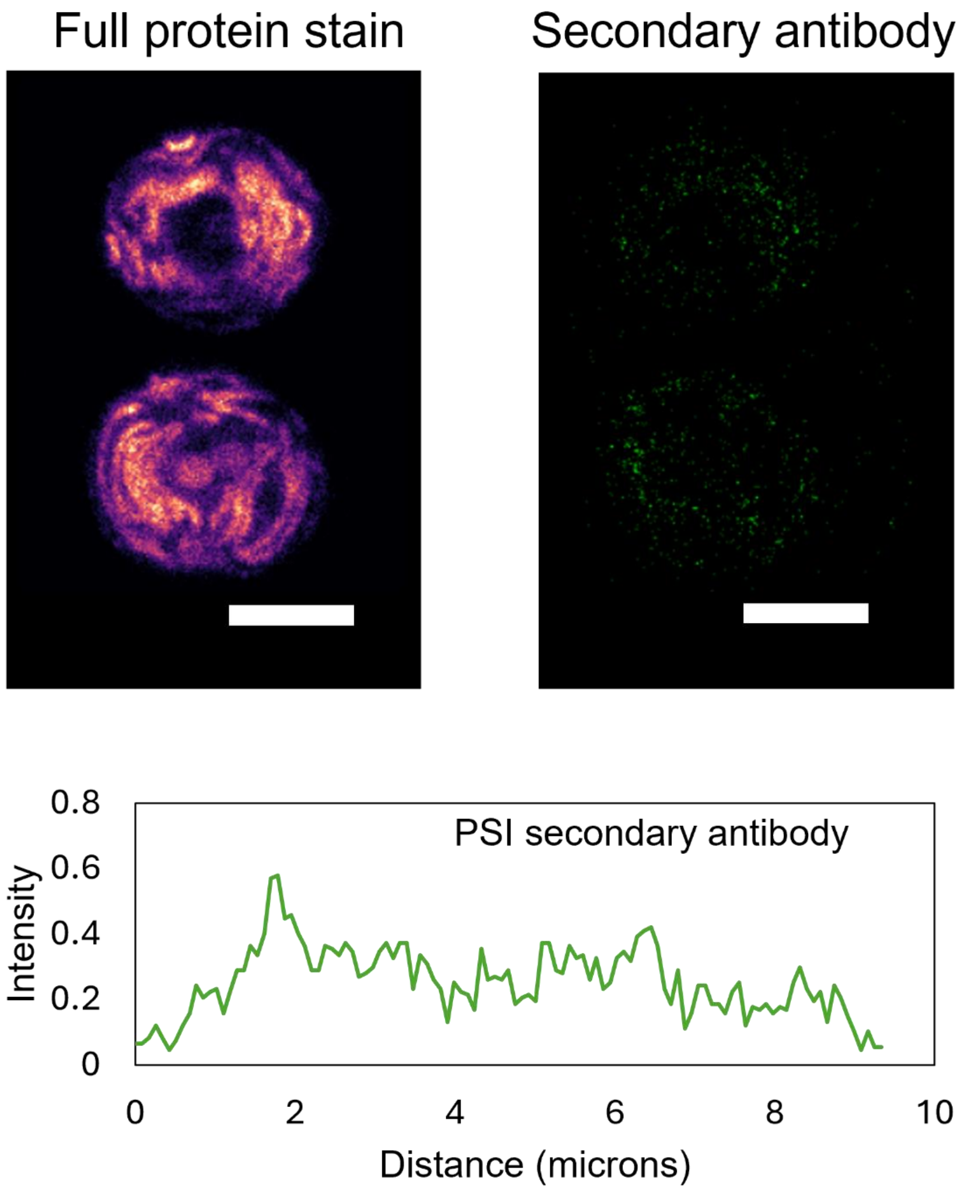


Supplemental Figure 2 Example of only secondary antibody staining as negative control. Full protein staining (ATTO-647N-NHS) on the left and secondary antibody staining (Alexa-488 nm) on the right. In the bottom panel, the average pixel intensity on a scale from 0 to 255 of the secondary antibody image is shown, from top to bottom.

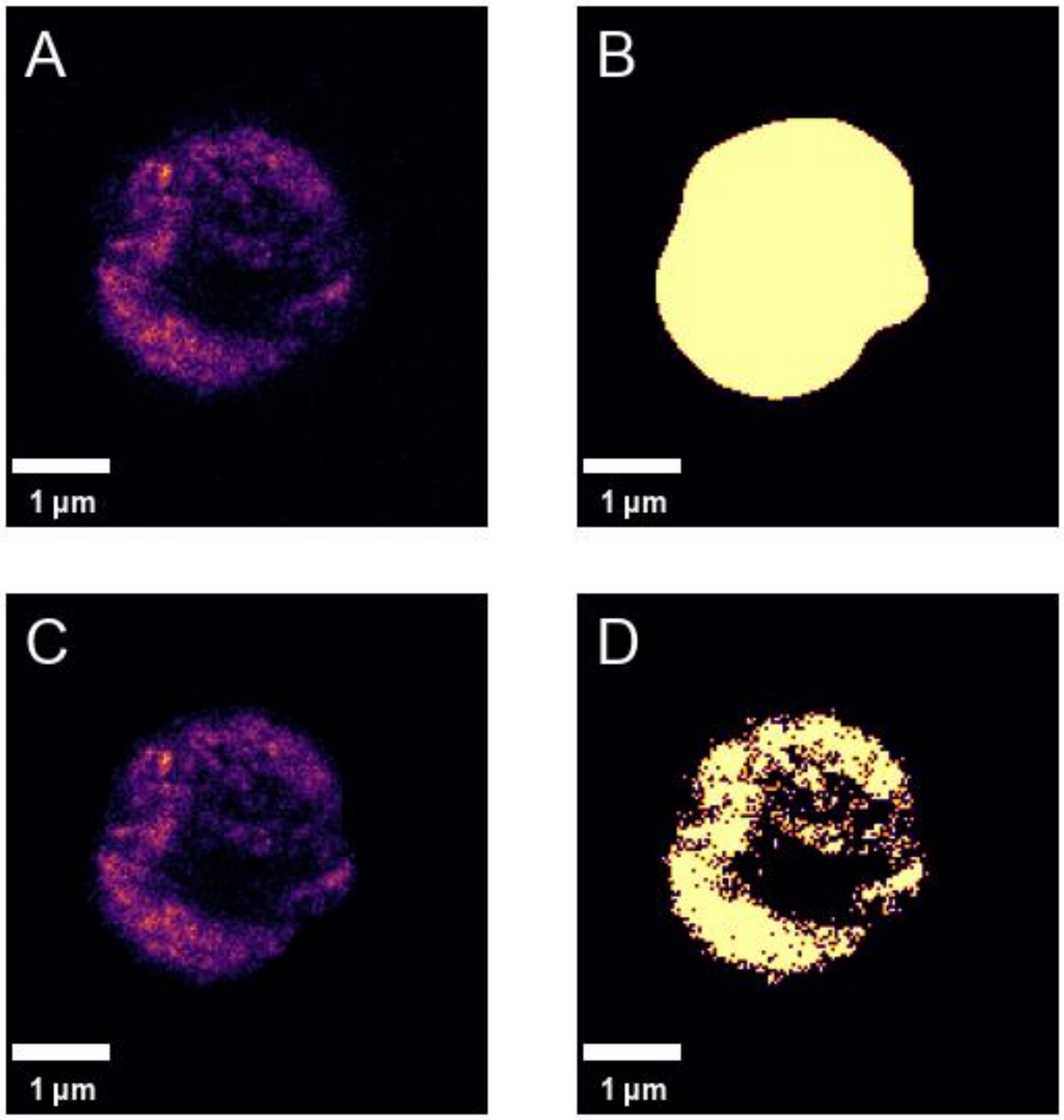


Supplemental Figure 3 Example of image treatment in order to determine the PCC and MCC values. A) Original image in the inferno color scheme. B) Original image, binarized with the Otsu filter. Pixel intensity is either 0 or 1. C) Multiplication of image A and B, results in the usual intensity inside the masked area, and intensity 0 outside the area of interest. D) Binarized image C with an Otsu filter. PCC was determined from image C from two different channels. MCC can be determined from image D from one channel and image C from another.

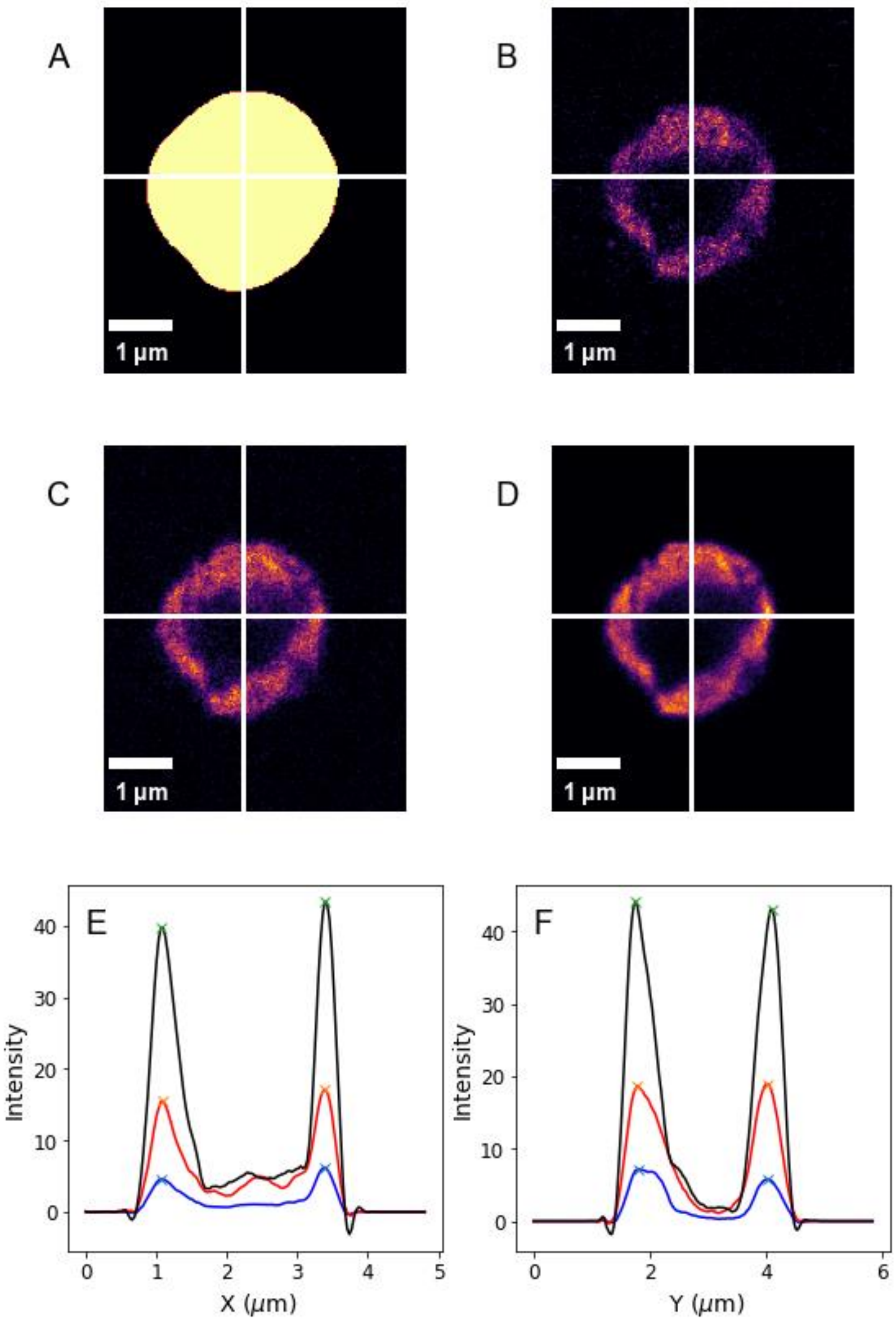


Supplemental Figure 4 Example of image analysis to determine the peak position of the antibodies on the thylakoid membrane. A) Micrograph binarized by an Otsu filter. The rows and columns of the image with the largest number of masked pixels are determined as the center of the cell. B, C, D) Images of PSI, PSII and phycocyanin respectively with the profile lines indicated in white. E, F) Smoothed profiles along the X and Y axis, respectively. Peak positions of phycocyanin (black) are indicated by green marks, PSII (red) with orange ones and PSI (blue) with light blue ones. The mean positions were determined and with that the difference of each channel peak from the mean. These differences were multiplied with the pixel size corrected for the expansion factor and plotted in figure 3B.

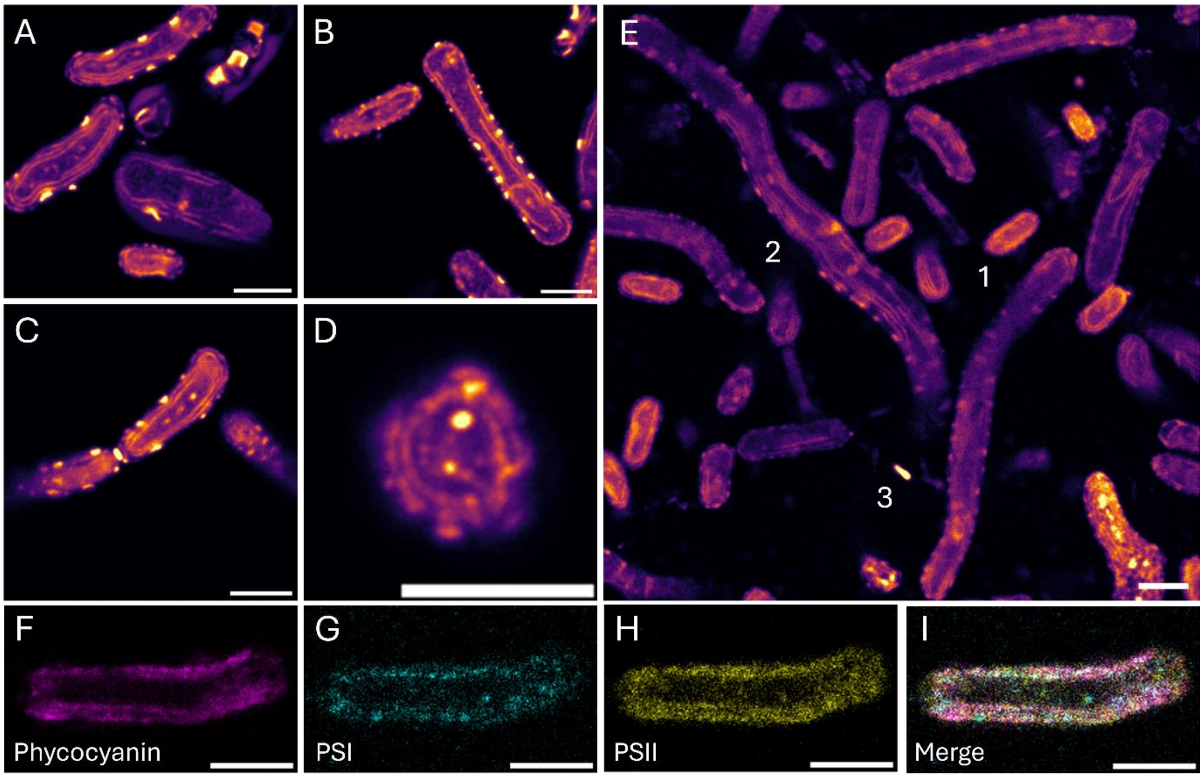


Supplemental Figure 5 Micrographs of ~5 times expanded *Synechococcus elongatus PCC 7942*. A-D) Expanded *Synechococcus* cells stained with all-protein staining. Side-views (A, B and C) and top-views (D) were found. E) anti-C-phycocyanin stained cells show the diversity in cell-shapes occasionally observed when expanding *Synechococcus*. At 1 a cell with the expected size based on the expansion factor, at 2 a cell that is about 6 times longer and a bit wider. The cell is not a rigid rod, but rather forms a wave shape. At 3 an unexpanded cell that is still stained by anti-C-phycocyanin. F-I) Expanded *Synechococcus* cells stained with anti-C-phycocyanin, anti-PsaC or anti-PsbA, and the merge of the three channels. Both PSI and PSII displayed a low intensity staining with maximum intensity of 7 or 15 respectively in an 8=bit image. Scalebars represent 1 µm in pre-expanded dimensions.
